# Can we trust subthalamic local field potentials? Geometric and dynamical constraints on the interpretation of extracellular recordings

**DOI:** 10.64898/2026.07.08.737284

**Authors:** F. Fattorini, T.V. Ness, L.G. Amato, A. Mazzoni, N. Meneghetti

## Abstract

Local field potentials (LFPs) are widely interpreted as readouts of population synaptic activity, an assumption derived almost entirely from cortical recordings. Whether these principles extend to subcortical structures remains unclear. We address this question in the subthalamic nucleus (STN), where LFPs are routinely recorded and used to guide adaptive deep brain stimulation for Parkinson’s disease, using a biophysically detailed population model benchmarked against patient microelectrode recordings. As in the cortex, STN extracellular potentials were dominated by synaptic currents. Differently from the cortex, however, LFPs could not be reliably predicted from these currents or other average population quantities. This dissociation arises from the STN symmetric neuronal morphology and lack of recurrent connectivity, which promote destructive interference among single-neuron contributions, decoupling the LFP from population-level dynamics. This decoupling was not absolute: pathological beta synchrony restored a robust synapse-LFP relationship by consistent underlying dynamics, while the aperiodic slope of the power spectral density tracked STN neuronal morphology, firing rate, and excitatory-inhibitory balance. Together, these findings challenge the prevailing view of LFPs as universal readouts of population activity. Our results show that the interpretability of extracellular signals depends critically on neuronal morphology and synchronization state, and provide a mechanistic framework for the use of STN LFPs as biomarkers in adaptive deep brain stimulation for Parkinson’s disease.

## Introduction

Extracellular potentials (ECPs) are among the most widely used electrophysiological signals in neuroscience^1^. They reflect neuronal transmembrane currents in the vicinity of the recording electrode, including contributions from spiking and synaptic activity^2,3^. Owing to their rich informative content, ECPs are both used in basic and clinical neuroscience as biomarkers of neurological disorders^4–6^.

The physiological mechanisms underlying ECP generation have been extensively investigated in cortical circuits^7–12^, where local field potentials (LFPs), i.e., low-pass filtered ECPs, primarily reflect synaptic inputs onto spatially aligned pyramidal neurons. In contrast, LFPs in subcortical structures remain poorly understood. This gap between cortical and subcortical electrophysiological features limits a mechanistic interpretation of experimental recordings and the development of computational models of subcortical networks.

These limitations are particularly relevant for the subthalamic nucleus (STN), a key structure in Parkinson’s disease (PD) and the primary target of deep brain stimulation (DBS)^13–15^. STN LFPs are routinely recorded to characterize pathological network dynamics and guide adaptive DBS strategies^16–18^. However, despite numerous studies linking LFP features to PD symptoms^14–17^, linking pathophysiological mechanisms with STN electrophysiological signals remains a major challenge. Computational modelling offers a powerful approach to address these questions^19,20^. However, most PD computational studies have focused on the mechanisms underlying pathological synchronization of firing dynamics inside the basal ganglia, and in particular within beta^21,22^ and gamma^23^ bands. The few computational studies that have explicitly incorporated neuronal morphology (a necessary component to accurately estimate ECPs) have primarily examined the effects of DBS within the STN^25,26^, rather than systematically investigating how morphology, synaptic inputs, and firing dynamics interact to shape ECPs.

Here, we address this issue leveraging a biophysical model of the STN incorporating detailed neuronal morphology, anatomically constrained synaptic connectivity, and realistic population architecture. The resulting ECPs were benchmarked against microelectrode recordings from PD patients. We show that synaptic currents contribute substantially to STN ECPs, as in the cortex. However, unlike cortical ECPs, STN ECPs cannot be reliably predicted from synaptic currents. We identify neuronal morphology and synaptic input dynamics as the main determinants of this dissociation, and we show that pathological beta oscillations can restore this coupling. Finally, we demonstrate that neuronal morphology and excitatory/inhibitory balance shape the STN spectral aperiodic slope.

Together, these results establish a mechanistic link between neuronal architecture, synaptic dynamics, and subthalamic extracellular signals, providing a foundation for the interpretation of STN recordings in health and disease.

## Results

### Validation of synaptic parameters against biological constraints

Building upon a previously developed multicompartmental model of an STN neuron^27,28^, we tuned synaptic features^25^ (Fig. 1A) to match literature-reported findings. We placed peri-somatic cortical (i.e., excitatory) and dendritic-targeting pallidal (i.e., inhibitory) synapses and performed voltage-clamp simulations in response to individual input spikes. We analyzed the amplitude of the simulated post-synaptic currents and identified the range of synaptic conductances reproducing experimentally reported values (excitatory currents: 105 ± 5 pA^29^; inhibitory conductances: 0.24 ± 0.06 nS^30^, gray bar in Fig. 1B). Excitatory synapses required larger conductances and showed larger variability, yet produced amplitudes comparable in amplitude to inhibitory synapses, indicating a higher efficacy of peri-somatic inhibition. Within the selected ranges, we next investigated which combinations of excitatory and inhibitory conductances reproduced the firing statistics of STN neurons recorded in vivo^31–33^. Simulations used literature-based estimates of synaptic numbers and input firing rates for cortical and pallidal populations. This analysis revealed a well-defined region of parameter space in which the model produced physiological firing rates (Fig. 1C, black dots) with mean value between 30 to 40 spikes/s and coefficients of variation close to 1.0. Outside this region, the model exhibited two distinct failure regimes. Insufficient excitatory drive produced low-rate or absent firing (Fig. 1C). Conversely, excessive excitation produced sustained depolarization and loss of physiological firing (Fig. 1C white region). Based on these constraints, we selected inhibitory and excitatory synaptic conductances of 0.28 nS and 15 nS, respectively (dashed lines in Fig. 1B, C; Methods), which reproduced the experimentally observed firing properties. Under these synaptic parameters, the model transitioned from the regular spontaneous firing observed in the absence of synaptic inputs (Fig. 1D top) to an irregular firing regime consistent with in vivo STN activity (Fig. 1D bottom).

**Fig. 1.**
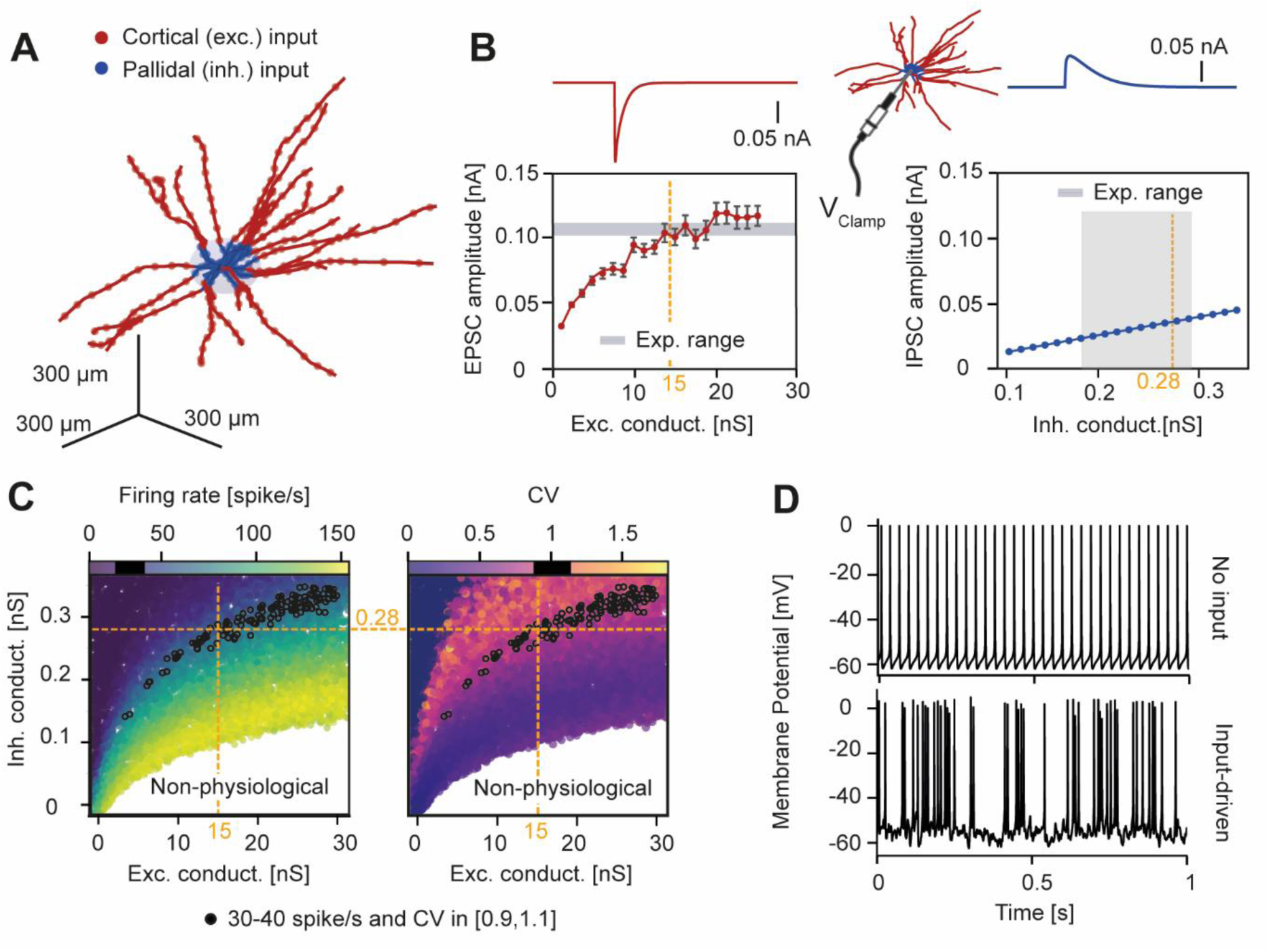
Biological constraints on single-neuron synaptic dynamics. A) Morphological model of subthalamic nucleus neuron, receiving perisomatic (blue) pallidal inputs and distal (red) cortical inputs. B) Amplitude of excitatory (left) and inhibitory (right) evoked post-synaptic currents (EPSC) in a simulated voltage-clamp experiment as a function of synaptic conductances. Experimental ranges taken from literature^29,30^ are represented in gray. C) Mean firing rate (left) and coefficient of variation (CV, right) across pallidal (inh.) and cortical (exc.) synaptic conductances. Black dots indicate simulations reproducing experimentally observed firing statistics^31–33^. White regions correspond to non-physiological firing regimes. The conductance values used in subsequent simulations are indicated by dashed orange lines. D) Representative STN membrane potentials without (top) and with (bottom) synaptic inputs.

### Role of synaptic inputs in the STN extracellular potentials

With single-neuron dynamics and synaptic inputs established, we evaluated whether an STN population with a realistic architecture could reproduce experimentally observed extracellular signals. We placed STN neurons neurons within a 600 µm-radius sphere with uniform density^34^ and random orientations (Fig. 2A, Methods and Suppl. Fig. 1). Using a well-established forward-modeling framework that combines multicompartmental neuronal modeling and volume conduction theory^35,36^, we computed ECPs at the population center and compared their power spectral density (PSD) with the one computed from STN microelectrode recordings of PD patients^37^ (Fig. 2B; see Methods).

**Fig. 2.**
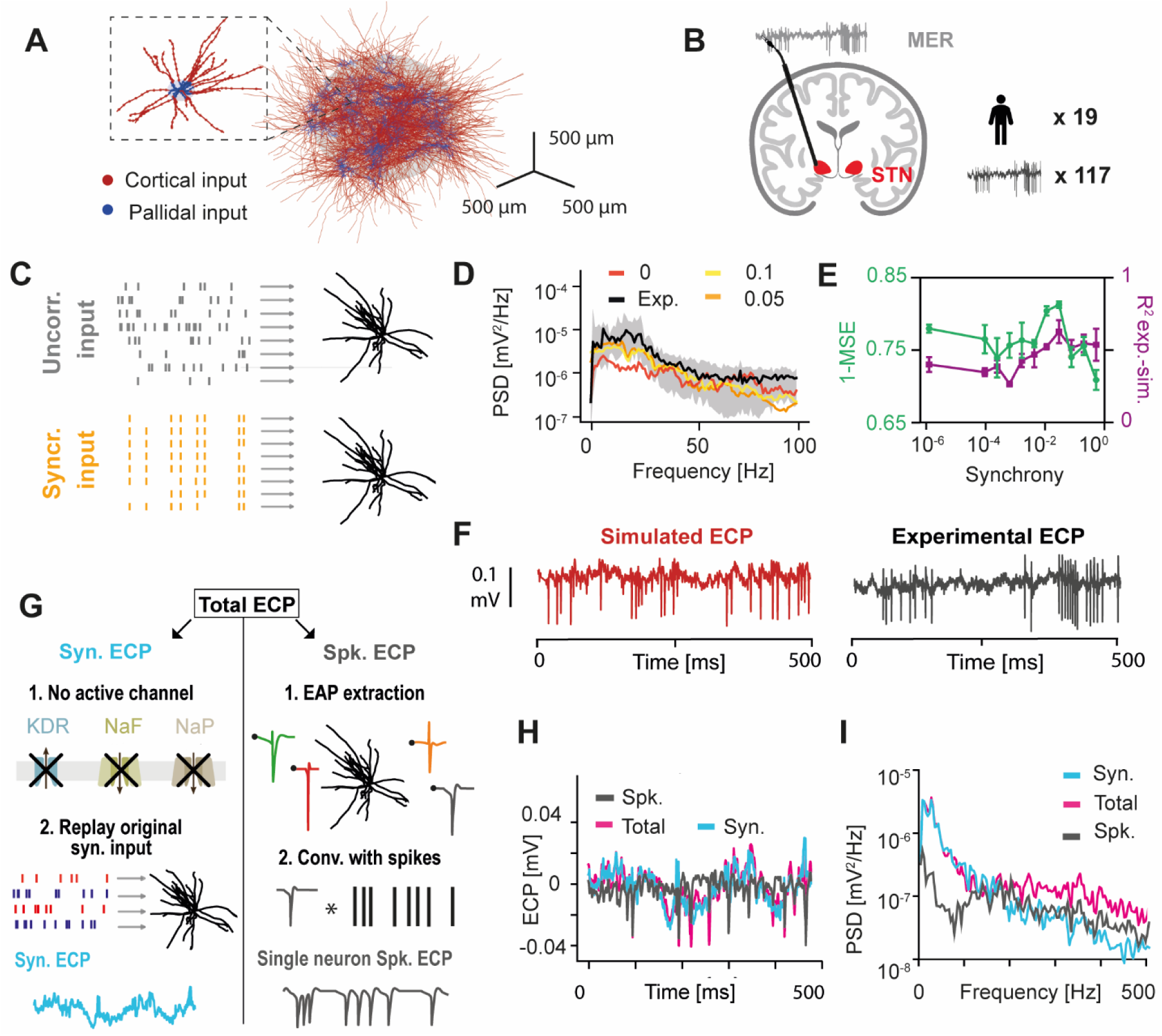
Simulated extracellular potentials reproduce experimental recordings and separate synaptic and spiking contributions. A) STN population model used for ECPs simulations. B) STN microelectrode recordings (MERs)^37^ used for comparison with simulations (19 patients and 117 recordings). C) Representative scheme illustrating uncorrelated (top) and synchronized (bottom) synaptic inputs to a STN neuron. D) Median power spectral densities (PSDs) of experimental recordings and simulations obtained at different synchrony levels. E) Mean squared error (MSE) and coefficient of determination (R^2^) between simulated and experimental PSDs across input synchrony (top). F) Representative simulated (left) and experimental (right) ECPs show comparable amplitudes. G) The schematic illustrates the computational pipeline used to decompose the simulated ECP into synaptic and spiking components. H) Total ECPs with their synaptic and spiking contributions. I) PSDs of the synaptic, spiking, and total ECPs.

We then varied the degree of synaptic synchrony by increasing the probability of coincident synaptic events delivered to individual neurons (Fig. 2C). In the absence of synchrony, the simulated ECP showed markedly lower spectral power than the intraoperative STN recordings from PD patients (Fig. 2D), indicating that independent synaptic drive alone is insufficient to reproduce the experimental spectrum. Introducing synaptic synchrony improved agreement with the recordings only within a restricted range of synchrony (Fig. 2F), whereas higher levels of synchrony progressively degraded the fit quality. We therefore set the input synchrony to 0.05, which maximized the goodness of fit with experimental PSDs (MSE=0.180 ± 0.008, R² = 0.66 ± 0.14). Importantly, the model failed to capture the beta-band PSD observed in the recordings regardless of synchrony (Fig. 2D), consistent with the absence of pathological beta hypersynchrony in the current model. Nevertheless, the simulated ECP amplitudes closely matched those of the experimental microelectrode traces, as shown in Fig. 2F.

We then investigated the relative contributions of synaptic and spiking activity to the modeled ECPs. To isolate synaptic contributions, we constructed a passive version of the STN population in which neurons received the same synaptic inputs as the full model but lacked active ion channels required for action potential generation. This allowed us to compute ECPs driven exclusively by synaptic currents (Methods; Fig. 2G, left). To assess the contribution of action potentials, we reconstructed the spiking component of the ECP by convolving, for each neuron, its extracellular action potential waveform with the spike times, and then summing the contributions across neurons (Fig. 2G, right; Methods). These approaches allowed us to disentangle synaptic and spiking contributions to the STN ECP under the same underlying dynamics (Fig. 2H). We found that synaptic activity accounted for most of the variance in the full ECP (R² = 0.66), whereas spiking activity provided a smaller contribution (R² = 0.17). Accordingly, the low-frequency component of the ECP, which contains most of the spectral power, was dominated by synaptic currents, whereas action potential contributions became comparable only above ∼150 Hz (Fig. 2I). This pattern is consistent with cortical recordings^7^, although the crossover frequency in the cortex is substantially higher (>300–400 Hz).

### Symmetric morphology and uncorrelated activity limit STN LFP reconstruction

In cortical circuits, LFPs are largely generated by synaptic activity^1,7^. Importantly, this relationship can be inverted, allowing LFPs to be reconstructed from a linear combination of excitatory and inhibitory population synaptic currents^8^. Having shown that STN ECPs are likewise dominated by synaptic currents (Fig. 2I), we next asked whether the same synaptic-to-ECP relationship extends to the STN (Fig. 3A). We first validated this approach in a cortical network composed of simplified ball-and-stick excitatory and inhibitory neurons (Fig. 3A, right; Methods). Consistent with previous studies^8^, cortical LFPs were accurately reconstructed from the sum of the absolute excitatory and inhibitory synaptic currents (R² = 0.94; Fig. 3B, right). Applying the same approximation to the STN model yielded almost no correspondence between synaptic currents and the ground-truth LFP (R² = 0.03; Fig. 3B, left), indicating that STN LFPs cannot be linearly reconstructed from population synaptic activity. Similar results were obtained using population firing rates, average membrane potentials (Supplementary Fig. 2A) or kernel-based approaches^9,38^ (Supplementary Fig. 2B-C), suggesting that STN LFPs cannot be reliably inferred from standard population-level measures of network activity. The divergence between cortical and STN LFP generation became evident as larger fractions of the neuronal population were included in the extracellular signal reconstruction (Fig. 3C). In the cortical model, reconstruction accuracy improved with additional neurons, consistent with population averaging reducing single-neuron variability. In contrast, reconstruction performance deteriorated in the STN as more neurons were included, indicating that population summation does not average out single-cell variability in this structure. This effect became evident when neuronal dynamics were preserved but somatic positions were randomized (Fig. 3D). Cortical excitatory LFPs were largely unchanged, with correlations between original and permuted signals remaining near 1. In contrast, STN LFPs were highly sensitive to neuronal arrangement, with correlations approaching 0 after permutation. Cortical inhibitory populations showed an intermediate response (R² = 0.44). Together, these results indicate that STN extracellular signals remain strongly shaped by randomness in the spatial organization, whereas cortical LFPs primarily reflect population-level activity.

**Fig. 3.**
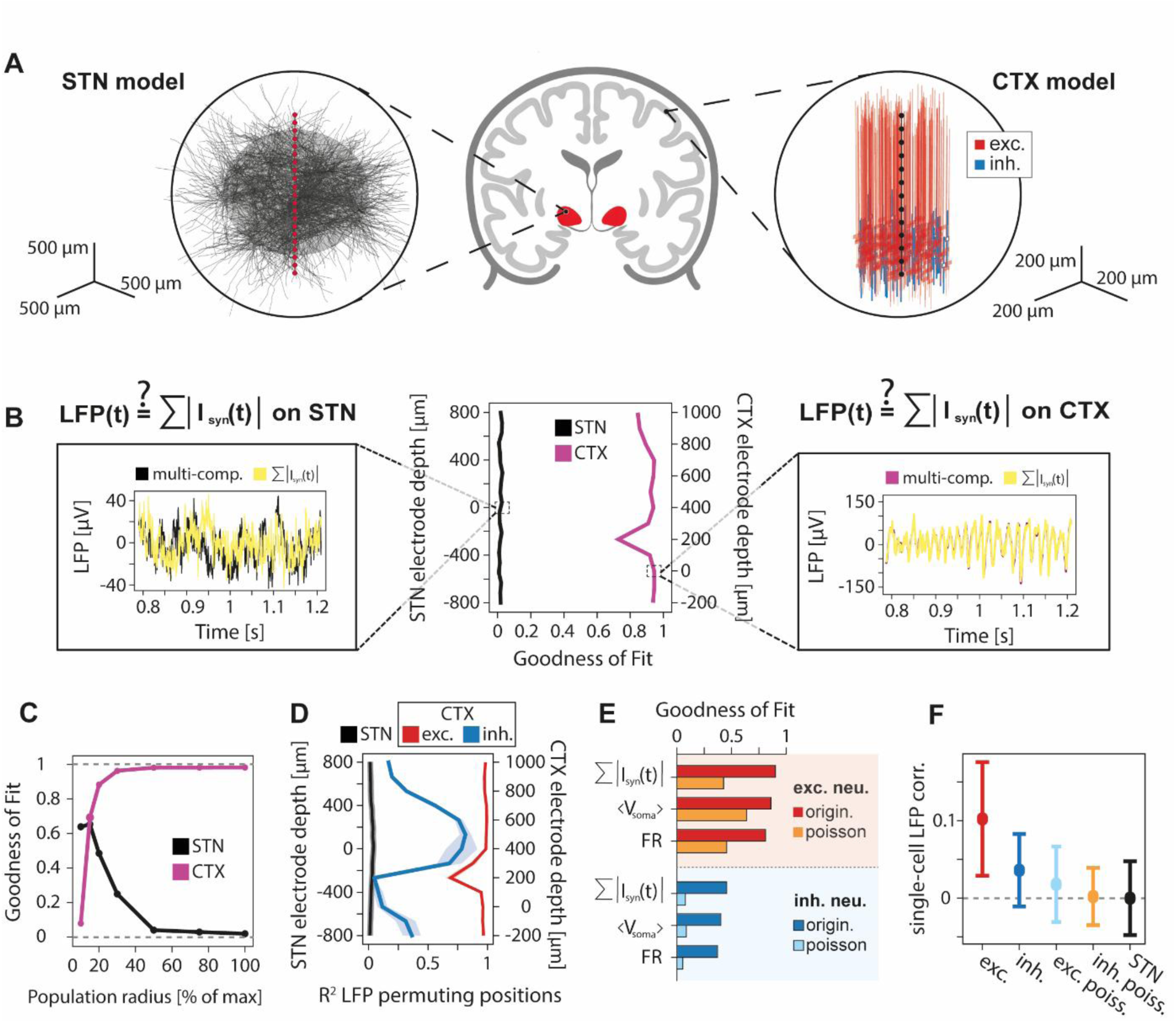
Weakly correlated single-neuron contributions limit population-level reconstruction of STN LFPs. A) Schematic of the STN (left) and ball-and-stick cortical networks (right). B) Goodness-of-fit (R²) of synaptic-current-based LFP reconstruction across recording depths for STN and cortical network. Insets show reconstructed and ground-truth LFPs. C) Coefficient of determination (R²) of synaptic-current-based LFP approximation across population radius in STN and cortical networks. R² between original and somatic position permutated LFPs, across recording depths, for STN and cortical excitatory and inhibitory populations. E) R² between ground-truth LFPs and population-level descriptors, including synaptic current sum (I_syn_), mean somatic membrane potential (⟨V_soma_⟩), and firing rate (FR). Results are shown for cortical excitatory and inhibitory populations under original connectivity and under Poisson-driven inputs. F) Pairwise correlation coefficients between LFP signals generated by individual neurons, in cortical and STN populations.

This intermediate behavior of cortical inhibitory populations suggested that neuronal symmetry, a feature that makes inhibitory neurons more STN-like than cortical excitatory neurons, can only partially account for the observations in the STN. We therefore tested the role of recurrent connectivity, a hallmark of cortical circuits that is absent in the feedforward STN network, by replacing recurrent cortical inputs with Poisson spike trains matched for mean firing rate, thereby disrupting coordinated synaptic activity generated by local network interactions. This manipulation substantially reduced the goodness-of-fit of the synaptic LFP approximation for excitatory neurons, lowering R² from 0.94 to 0.47 (Fig. 3E). The effect was even stronger for inhibitory neurons, where R² decreased from 0.45 to 0.09, reaching values comparable to those observed in the STN. Similar effects were obtained when reconstructing the LFP from other population-level descriptors, including average somatic membrane potential and firing rate (Fig. 3E).

These results suggest that LFP interpretability is primarily determined by the correlation structure of single-neuron contributions to the population field, which is influenced by neuronal morphology, spatial arrangement, and synaptic input correlations. Consistent with this, pairwise correlations between single-cell LFP traces were high in cortical excitatory populations, reduced in inhibitory populations, and further decreased after replacing recurrent interactions with Poisson inputs (Fig. 3F). In both Poisson-driven cortical networks and the STN, correlations approached zero, indicating that the ECP reflects the summation of largely uncorrelated neuronal contributions. Even when identical inputs were delivered to all STN neurons at differing synaptic locations, the resulting LFPs remained weakly correlated (r = 0.008 ± 0.075), suggesting that the symmetric morphology and placement of synapses are sufficient to strongly decorrelate the single LFPs.

### Pathological beta synchrony restores LFP interpretability

The results described above indicate that STN LFPs are strongly shaped by cancellation effects, weakly related to standard population descriptors, and highly sensitive to recording geometry (Fig. 3), suggesting a noise-dominated signal with limited information content. This contrasts with extensive experimental and clinical evidence showing that STN LFPs exhibit structured, disease-related spectral features, including beta-band oscillations^18,39^ and the aperiodic slope^40^ of the power spectrum, both well-established biomarkers in Parkinson’s disease. We therefore asked whether structured network synchronization could restore the information content of STN LFPs.

To address this question, we introduced beta-oscillatory cortical and pallidal inputs into the model, independently varying their amplitudes and relative phase shift (Fig. 3A). Model parameters were constrained using the same STN microelectrode^37^ recordings employed in previous analyses by matching spike-triggered beta-band (13–30 Hz) LFPs. The best-fitting solutions consistently exhibited a cortical–pallidal phase shift between 0 and π/2 (≈11 ms; Fig. 4A), together with a dominant cortical oscillatory drive (A_exc_ = 3 spike/s, A_inh_ = 7.5 spike/s). The corresponding best-fitting solution accurately reproduced the experimental LFPs (Fig. 4B–C), indicating that beta-band spike–field coupling is primarily governed by cortical oscillatory input and its phase relationship with pallidal activity.

**Fig. 4.**
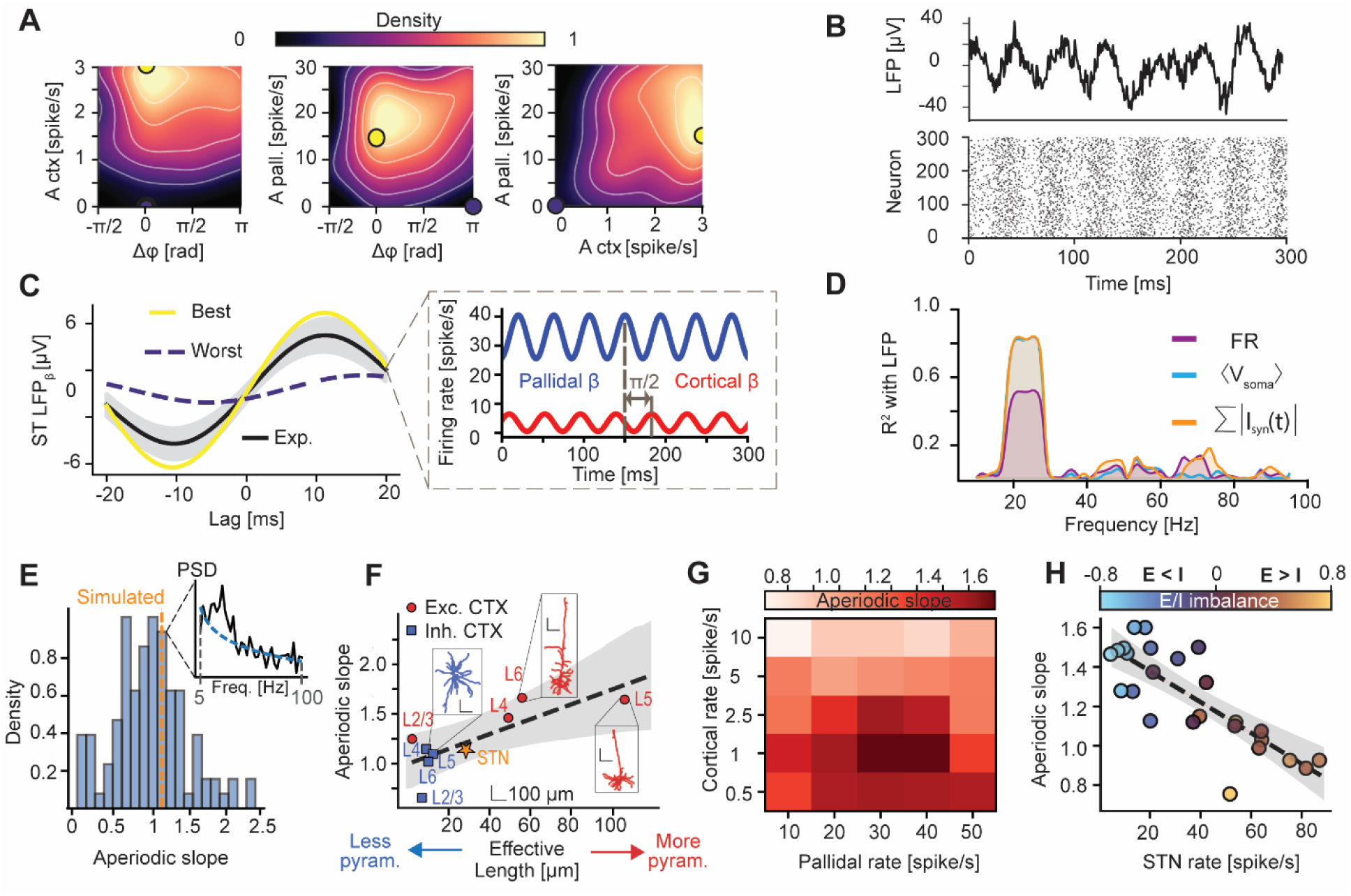
Beta synchronization and aperiodic components capture informative features of LFP signals. A) Distributions of model parameter (A_ctx_, A_pall._, and Δφ, see Methods) combinations that best reproduced the spike-aligned beta-filtered LFP across experiments. Yellow and purple dots indicate the best- and worst-matching parameter sets, respectively. B) Representative simulated LFP (top) and raster plot (bottom) for the optimal beta oscillation parameters. C) Left: experimental spike-aligned LFPs (black) and corresponding best (yellow) and worst-matching (purple) simulations. Right: schematic of cortical and pallidal input currents in the best-matching configuration. D) Band-wise coefficient of variation R^2^ between population variables (Firing rate FR, mean membrane potential ⟨V_soma_⟩, and sum of synaptic currents I_syn_) and simulated LFPs. E) Distribution of experimental LFP aperiodic slopes (5–100 Hz) and simulated value. Top-right panel shows an example of aperiodic component extraction from experimental PSD. F) Relationship between neuronal effective length (i.e., morphology asymmetry, see Methods), and the LFP aperiodic slope in the Allen Institute V1 model. Excitatory and inhibitory neurons are shown in red and blue squares, respectively. The dashed line indicates the linear fit, and the shaded region its 95% confidence interval. The orange star indicates the STN neuron. G) Simulated LFP aperiodic slope as a function of cortical and pallidal input firing rates. H) Simulated LFP aperiodic slope as a function of STN firing rate. Color indicates excitation-to-inhibition (E/I) imbalance; the dashed line denotes the linear regression.

The introduction of beta oscillations qualitatively transformed the relationship between population activity and the LFP. Whereas synaptic currents and membrane potentials poorly predicted STN LFP in the absence of beta (Fig. 3), both quantities accurately followed beta-band LFP fluctuations under oscillatory conditions (Fig. 4D). Thus, pathological synchronization restores a low-dimensional relationship between population activity and the STN LFP that is otherwise absent from the asynchronous regime.

### STN aperiodic slope reflects neuronal morphology, E/I balance, and average STN firing rate

Having characterized the rhythmic components of the STN LFP, we next investigated its aperiodic spectral component. The aperiodic exponent simulated in out model (1.13) fell within the range observed in the dataset of experimental MER recordings (Fig. 4E, mean ± STD=1.00 ± 0.49) and was substantially lower than values typically reported for cortical recordings (approximately 2.5 - 3.0^41^). To investigate the origin of this difference, we examined how neuronal morphology shapes the PSD aperiodic slope using morphologically diverse neurons from the Allen Institute visual cortex database^42^ (see Methods). Across cell types, the aperiodic exponent increased monotonically with morphological asymmetry (r = 0.78, p=0.02 Fig. 4E): neurons with elongated, pyramidal-like morphologies produced steeper spectral slopes, whereas more symmetric inhibitory interneurons exhibited shallower exponents. STN neurons fell on the same relationship, consistent with their relatively symmetric morphology. Interestingly, applying this relationship to a reconstructed human layer 5 pyramidal neuron yielded a predicted PSD slope of 2.7, in quantitative agreement with human cortical recordings^41^. These results indicate that the aperiodic exponent is strongly constrained by neuronal morphology and that a simple morphological relationship can account for regional differences in LFP spectral slopes. While neuronal morphology explained the lower aperiodic slopes of the STN relative to cortex, STN network dynamics also contributed to shaping the aperiodic exponent. The PSD slope was also strongly related to the STN mean firing rate (r = −0.81, p << 0.001; Fig. 4F) and to the balance between excitatory and inhibitory inputs, with stronger inhibition associated with steeper slopes (r = −0.81, p << 0.001; Fig. 4F). The aperiodic slope therefore inherits STN sensitivity to firing rate through its dependence on the E/I drive ratio, linking an arrhythmic spectral feature of the LFPs to the spiking regime of the network. Together, these findings show that STN LFPs, despite their limited relations with population quantities under asynchronous conditions, can retain information about network dynamics: the emergence of beta synchronization (observed in PD) restores a direct link with synaptic activity, whereas the aperiodic slope reflects the underlying firing rate and excitatory–inhibitory balance.

## Discussion

Local field potentials are widely considered indicators of population-level neural activity, but their interpretation has largely been derived from cortical regions. Our results show that these assumptions do not generalize to the subthalamic nucleus. Using a biophysically detailed STN model validated against microelectrode recordings from Parkinson’s disease patients, we demonstrate that, although STN ECPs are primarily generated by synaptic currents, they cannot be reliably reconstructed from those currents. This dissociation differs fundamentally from cortical regions and arises from two key properties: the symmetric morphology of STN neurons and the absence of recurrent connectivity, which reduces correlated synaptic activity. Together, these features promote destructive interference when summing single-neuron contribution to field potentials, making STN LFPs highly sensitive recording position. Nevertheless, STN LFPs can retain meaningful information about network dynamics. In particular, the aperiodic spectral slope reflects the balance between excitatory and inhibitory drive and the resulting firing regime, suggesting a potential biomarker of network state. Moreover, under conditions of (pathological) beta synchrony, the relationship between synaptic currents and the LFP is substantially strengthened.

Together, these results reveal a fundamental divergence in how extracellular signals reflect population activity across cortical and subcortical circuits, calling for a circuit-specific framework for interpreting field potentials.

### Comparison with previous modeling studies

Previous computational studies of STN LFPs have typically extended cortical models, assuming a direct, geometry-independent mapping between population synaptic currents and LFPs, modeled as their summed network activity^43–45^. Our results challenge this assumption by showing that neuronal morphology, spatial organization, and recording positioning strongly influence STN ECPs. Other studies have incorporated detailed STN morphology and detailed DBS electrode modeling^25,26,46,47^. In comparison, our model focuses on a more systematic characterization of synaptic mechanisms, including dynamics, conductance values, and synapse number per neuron, while exploring how different cortical and pallidal input configurations shape STN extracellular signals. In this sense, our work is complementary to previous approaches: whereas prior studies have primarily addressed electrode properties and the effects of DBS, we investigate the relationship between spiking activity, synaptic dynamics, and extracellular fields in the STN. Importantly, our results delineate conditions under which STN LFPs do not reliably reflect underlying population synaptic activity, providing a theoretical framework to guide the interpretation of STN recordings.

### Synaptic and spiking contributions to STN LFPs

In cortical regions, action potentials contribute little below approximately 300–500 Hz, allowing low-frequency ECPs (i.e., LFPs) to be primarily determined by synaptic signals^1,48,49^. In contrast, our simulations, combined with a novel method to properly disentangle spiking and synaptic contributions, show that spiking activity begins to dominate STN extracellular potentials above roughly 150 Hz. Consequently, broadband and high-gamma analyses in the STN likely contain substantial spiking contributions and should not be interpreted using cortical conventions^48,50^.

This difference stems from neuronal morphology and spatial architecture. Cortical pyramidal neurons possess elongated apical dendritic arbors that generate strong, spatially aligned dipoles, amplifying the effect of synaptic currents on extracellular field potentials^10,51^. The STN lacks this architecture^28^. The compact and symmetric morphology of its neurons reduces synaptic dipole strength and inter-neuronal alignment, thereby weakening the overall contribution of synaptic currents and reducing their spectral separation from spike-related activity.

### STN LFPs do not directly reflect population synaptic activity

Standard population descriptors that successfully approximate cortical LFPs^8,11,52,53^, including synaptic currents, firing rates, and average membrane potentials, fail to capture STN LFPs. Our results suggest that this limitation arises from how individual neuronal contributions combine to generate the population field. In cortical circuits, correlated synaptic inputs and aligned current dipoles promote constructive interference, establishing a robust relationship between extracellular signals and population-level variables. As increasingly larger neuronal populations contribute to the field, the correspondence between synaptic currents and LFPs improves.

In the STN, however, sparse recurrent connectivity and heterogeneous cortical and pallidal inputs^54,55^ produce weak correlations between the extracellular contributions of individual neurons. Their summation therefore promotes cancellation rather than amplification, preventing reliable recovery of the underlying population activity from the LFP.

Additionally, our simulations reveal a strong dependence of STN LFPs on electrodes placement. Identical population activity can generate markedly different extracellular signals depending on the spatial arrangement of neurons relative to the recording contact. This observation has important implications for DBS, where electrode position varies both across patients and across contacts on the same lead^56,57^. Consequently, differences in LFP amplitude or spectral features may reflect recording geometry rather than underlying physiological differences, warranting caution when interpreting these signals in adaptive DBS applications^18,58^.

### Beta oscillations and aperiodic slope restore interpretability of local field potentials

Although asynchronous STN activity produces cancellation-prone signals, beta oscillations fundamentally alter this scenario. By introducing correlated synaptic drive, beta-oscillating inputs restore a strong linear relationship between synaptic currents and the extracellular signal. Under these conditions, LFPs provide a faithful readout of synchronized subthreshold dynamics.

The aperiodic slope of the STN power spectrum has also emerged as a clinically relevant biomarker in PD^40,59,60^. Our results provide a mechanistic explanation for both its baseline value and its state dependence. The relatively shallower STN slope with respect to cortical recordings^41^ arises directly from neuronal morphology and can be predicted from the degree of morphological asymmetry of the neurons generating the field potentials. In addition, we found the slope to reflect the balance between excitatory and inhibitory drive and to be linearly related to the mean firing rate of the STN, consistent with previous experimental observations ^61^. In our model, increasing excitatory inputs flattens the slope, whereas stronger inhibition steepens it.

### Conclusions

These findings argue against a universal interpretation of LFP. The relationship between synaptic activity and extracellular fields critically depends on neuronal morphology, synchrony, and recording geometry. In cortex, these factors support a faithful representation of synaptic dynamics. In the STN, they instead lead to cancellation, strong geometry dependence, and a weak and indirect relationship to population activity. Consequently, STN LFPs should not be interpreted as a direct readout of neuronal population activity in the same sense as cortical LFPs, but rather as signals whose meaning is strongly context dependent. This redefines how subcortical extracellular signals should be interpreted and sets constraints for their uses as adaptive DBS biomarkers.

## Methods

### STN single neuron

To construct the subthalamic nucleus population model, we first developed a biophysically detailed single-neuron model (Fig. 1). We adopted a previously validated morphological and electrophysiological model of an STN neuron^28^ and extended it by incorporating excitatory cortical and inhibitory pallidal synaptic inputs. Synaptic inputs were spatially distributed according to anatomical evidence^25^: inhibitory pallidal synapses were placed proximally (<100 µm from the soma), whereas excitatory cortical synapses were distributed across more distal dendritic compartments (Fig. 1A). Pallidal synapses were modeled with rise and decay time constants of 1.1 ms and 7.8 ms, respectively ^62,63^, whereas cortical synapses exhibited faster kinetics, with a rise time of 0.1 ms and a decay time of 3 ms^64^. Reversal potentials were set to −80 mV for inhibitory synapses^65^ and 0 mV for excitatory synapses.

Synaptic conductances were calibrated to reproduce experimentally observed postsynaptic currents and physiologically realistic firing dynamics. To constrain synaptic efficacy, we performed in silico voltage-clamp experiments (Fig. 1B). The somatic membrane potential was held at −70 mV for excitatory inputs^30^ and −80 mV for inhibitory inputs^29^, consistent with the experimental protocols. For each synapse type, single synaptic events were delivered at 200 randomly sampled locations drawn from the prescribed spatial distributions, and the resulting postsynaptic currents were recorded. The amplitude distributions of excitatory and inhibitory currents were then compared with experimentally reported ranges^29,30^ (Fig. 1B). Because pallidal synapses exhibit strong short-term depression, inhibitory conductances were tuned using their experimental steady-state synaptic efficacy rather than peak transient amplitudes.

The model was subsequently extended to include full synaptic populations. The number of pallidal afferents was set to 883 based on anatomical estimates^66^ whereas cortical inputs were included using an 8:3 ratio relative to pallidal inputs, consistent with previous computational study^67^. Pallidal neurons were modeled as Poisson spike generators firing at a mean rate of 33 Hz^68^, while cortical afferents fired at a mean rate of 3.5 spikes/s^69^ also with Poisson statistics^70–72^.

To identify parameter regimes yielding physiologically realistic STN dynamics, we performed a random search over excitatory and inhibitory synaptic conductances, considering 1000 single neuron simulations of 2 seconds each (Fig. 1C). For each parameter set, we quantified the mean firing rate and the coefficient of variation (CV) of inter-spike intervals. Physiological activity was defined as firing rates between 30–40 spikes/s and CV values between 0.9–1.1, consistent with electrophysiological recordings reported in previous studies^31–33^. Finally, to identify the best synaptic conductances, we selected the parameter pair located closest to the center of the physiological parameter region, defined as the region reproducing experimentally reported firing statistics (30–40 Hz firing rate and CV between 0.9 and 1.1) and satisfying the constraint of the voltage-clamp simulations (excitatory weights in [12, 20] nS and inhibitory weights in [0.18, 0.29] ms).

### STN population and local field potentials

For the STN population, we adopted a spherical distribution of neurons with a radius of 600 µm (Fig. 2A). This radius was selected a posteriori based on an analysis of the contribution of neurons at increasing distances from the electrode to the computed LFP. Specifically, we evaluated the approximation of the total LFP obtained by progressively increasing the radius of the neuronal sphere and compared it with a reference LFP computed using a fixed radius of 900 µm (Suppl. Fig. 1). The selected radius of 600 µm corresponds to the point beyond which the additional contribution of more distant neurons became negligible, as indicated by the stabilization of the LFP variability and the high coefficient of determination between the cumulative LFP and the reference LFP. Therefore, only neurons within this radius were included in the final LFP computation. Local field potentials were estimated at the center of the sphere using the LFPy library^36^, assuming a constant extracellular conductivity of 0.3 S/m. For Fig. 3B and 3D we considered linear electrodes equally spaced between −800 and 800 µm.

The effect of correlations among the inputs to individual neurons was also investigated. Correlations were introduced in the spike trains provided as input to a single neuron^73^ by increasing the degree of synchrony among spikes delivered to different synapses (Fig. 2C). A correlation value of zero produced fully independent Poisson spike trains in input to the same neurons, whereas higher values introduced shared “parent” spike events across multiple trains, resulting in temporally aligned activity.

The network was simulated for 2 seconds with an integration step of 2^-5^ ms. In all simulations, the first 100 ms of simulation were not considered to neglect the non-stationary state of the network.

### Microelectrode recordings in PD patients

To compare simulation results with experimental data, we used a previously published public dataset^74^. The dataset consisted of STN microelectrode recordings obtained from 19 patients with Parkinson’s disease (Fig. 2B), resulting in a total of 117 recordings of varying duration. To obtain a power spectral density representative of the experimental local field potential, we computed the power spectral density for each recording using Welch’s method (segment length: 2 s; sampling frequency: 10 kHz)) and then calculated the median spectrum across recordings (Fig. 2D). Before computing the power spectral densities, simulated signals and experimental ones were resampled with linear interpolation not the common sampling frequency of 1 KHz.

The comparison between simulated and experimental data was performed using the coefficient of determination R^2^ and the normalized mean square error (MSE) (Fig. D, top), defined as:

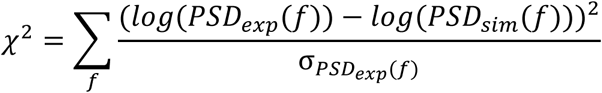

Where *PSD*_*exp*_(*f*) is the experimental power spectral density at frequency, *PSD*_*sim*_(*f*) is the simulated power spectral density at the same frequency and σ_*PSDexp*(*f*)_ denotes the standard deviation or uncertainty associated with the experimental PSD at frequency *f*.

### Decoupling of synaptic and spiking contributions to local field potentials

We disentangled the contributions of spiking and synaptic activity to the local field potential using two independent approaches (Fig. 2F). In both cases, we started from the complete network simulation and computed the LFP as the sum of the contributions from all individual neurons. To quantify the contribution of each component to the total LFP, we computed the coefficient of determination (R^2^, computed as squared correlation coefficient) between the reconstructed signals and the LFP obtained from the complete simulation. To isolate the contribution of synaptic activity (Fig. 2F, left), we replayed the same synaptic inputs from the complete simulation onto the same neuronal population, while removing the active membrane mechanisms responsible for action potential generation (i.e., fast sodium channels, persistent sodium channels, delayed rectifier potassium channels, Kv3.1 potassium channels, T-type calcium channels, high-voltage-activated calcium channels, small-conductance calcium-activated potassium channels, hyperpolarization-activated cation channels, intracellular calcium accumulation mechanisms). This procedure preserved synaptic currents while preventing neurons from producing spikes.

To estimate the contribution of spiking activity (Fig. 2F, right), we first extracted spike times from the full network simulation by thresholding the membrane potential at −10 mV. We then computed extracellular action potential templates for each neuron at the electrode position, preserving the original neuronal positions and rotations used in the full population simulation. Finally, we convolved each neuron’s spike train with its corresponding extracellular template to reconstruct the single-neuron contributions to the LFP.

### Cortical model

Cortical extracellular field potentials were simulated using the simplified ball-and-stick microcircuit model ^75^ (Fig. 3A). Excitatory and inhibitory neurons were represented by stylized multicompartmental models comprising a limited set of passive and active conductances distributed across somatic and dendritic compartments. Specifically, both cell types incorporated conductances derived from a previously validated biophysical model^76^, including the transient sodium current and the fast non-inactivating potassium current, which support spike generation, together with the hyperpolarization-activated cation current, which contributes to subthreshold dynamics. The network received external Poisson-distributed excitatory inputs and included recurrent synaptic connectivity both within and between excitatory and inhibitory populations. Detailed descriptions of channel kinetics, connectivity, and model parameters are provided in ^75^. Simulations generated two principal outputs: extracellular potential estimates and synaptic currents (Fig. 3B). Local field potentials were computed through LFPy^36^ in the center of the population (Fig. 3C, 3E and 3F) or with a linear electrode spacing from depth −200 µm to 1000 µm with respect to the center of the cortical population.

To assess the contribution of neuronal morphology to extracellular field generation, we separately quantified the extracellular signals generated by excitatory and inhibitory neuronal populations, as well as the total population signal. Synaptic currents were estimated as described in the section “Synaptic currents to approximate LFPs.”

To assess the contribution of recurrent synaptic connectivity to extracellular field potentials, we performed additional simulations in which all recurrent synaptic connections were removed (Fig. 3E, “Poiss”). Recurrent synaptic input was then replaced by independent Poisson-distributed spike trains, with rates set equal to the mean synaptic drive of the corresponding inputs in the original recurrent network.

### Synaptic currents to approximate local field potentials

To estimate synaptic currents from the multicompartmental simulations, we adopted the hybrid approach proposed by^52^. We stored, for each simulation, the somatic membrane potential of every neuron in the network, together with the timing of all synaptic inputs it received. For each synapse, the synaptic current was computed as the convolution of the presynaptic spike train with an alpha function describing the postsynaptic current time course, multiplied by the driving force given by the difference between the somatic membrane potential and the synaptic reversal potential. The total synaptic current of each neuron was then obtained by summing the contributions from all incoming synapses.

To estimate the LFP from the set of synaptic currents in the network (Fig. 3), we followed the procedure outlined in Mazzoni et al.^8^. Specifically, we approximated the LFP obtained from the multicompartment reference simulation as the sum across neurons of the absolute value of the total synaptic current:

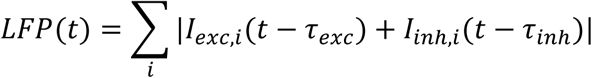

where I_exc_,_i_ and I_inh,i_ denote the excitatory and inhibitory synaptic currents impinging on neuron i, respectively, and τ_exc_ and τ_inh_ are time delays. These delays were varied from −20 to 20 ms in steps of 0.5 ms to maximize the R² between the estimated LFP and the reference LFP from the multicompartment simulation.

To test alternative approximation strategies, we also evaluated the goodness of fit between the reference LFP and the population-average somatic membrane potential, the instantaneous firing rate and a kernel-based linear prediction framework^9,75^ (Supplementary. Fig. 2).

### Beta oscillations

To incorporate beta oscillations into the model, we considered the relationship between spiking activity and LFP previously identified in the experimental dataset^37^. Rather than analyzing spike– LFP phase coupling directly, we computed spike-triggered averages of the beta-filtered (13-30 Hz) local field potential for each recording by averaging the LFP within a temporal window from −20 ms to 20 ms around each spike. Averaging these spike-aligned ECPs across recordings provided an estimate of the dominant beta oscillation frequency. We then fitted a sinusoidal function to the averaged trace, obtaining a characteristic frequency of 23 Hz. Then, beta-band activity was generated in the model by imposing sinusoidal temporal modulation on the firing rates of excitatory and inhibitory presynaptic populations. The instantaneous firing rates were defined as time-varying inhomogeneous Poisson processes with average rate of:

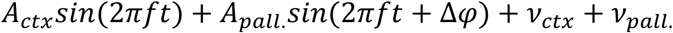

Where *A*_*ctx*_and *A*_*pall.*_ denote the amplitudes of the excitatory and inhibitory oscillatory inputs, respectively, *f* = 23 *Hz* frequency of the oscillation and *Δφ* defines the phase offset between the inhibitory and excitatory components. The terms *v*_*ctx*_ and *v*_*pall.*_ represent Poisson excitatory and inhibitory background activity, respectively. To avoid perfect synchrony across afferents, Gaussian temporal jitter was independently applied to each input neuron, with standard deviation σ=6.25 ms, following the methodology described by Lempka and McIntyre ^25^.

We compared the model with experimental data (Fig. 4A and 4C), performing a grid-search over values of *A*_*exc*_,*A*_*inh*_ and *Δφ* identifying for each recording the parameter combination that best reproduced the experimental spike-aligned LFP. The quality of the fit was quantified using a combination of the coefficient of determination and the mean squared error between simulated and experimental traces. This procedure produced a distribution of optimal parameter combinations across recordings. The most frequent parameter combination was then selected for the subsequent analysis. The marginal distributions shown in Fig. 4A were obtained by projecting the multidimensional parameter distribution onto each parameter independently and smoothing the resulting histograms using a kernel density estimator for visualization purposes.

Using these optimized parameters, we quantified the ability of different population-level variables to predict the LFP in specific frequency bands (Fig. 4D). Specifically, we evaluated firing rate, average excitatory synaptic input, average inhibitory synaptic input, and mean membrane potential. For each variable, both the predictor and the LFP were band-pass filtered in sliding frequency bands of 10 Hz width before computing prediction performance.

### Estimation of the aperiodic slope of LFP power spectral density

We estimated the spectral slopes in the 5–100 Hz range in the simulated and experimental recordings (Fig. 4E) by computing their power spectral densities and using the FOOOF method^77^ for the extraction of the aperiodic component of the spectrum (Fig. 4E, top-right inset).

To examine the relationship between neuronal morphology and the aperiodic slope of the LFP power spectrum, we analyzed extracellular potentials generated by a previously published large-scale model of mouse primary visual cortex (V1)^9^. The model comprises 230,924 neurons distributed across cortical layers 2/3, 4, 5, and 6, including 51,978 biophysically detailed multicompartment neurons arranged within a cylindrical core (800 μm diameter, 860 μm height) and 178,946 leaky integrate-and-fire point neurons forming a surrounding annular region (445 μm thickness) to minimize boundary effects. Full model architecture, biophysical parameters, and simulation details are provided in ^9^.

For each cortical layer, compound extracellular potentials were computed separately for excitatory and inhibitory populations by summing the extracellular contributions of all neurons belonging to each population. Signals were z-scored prior to spectral analysis to normalize for differences in population size. Power spectral densities were estimated using Welch’s method (segment length: 2 s; sampling frequency: 10 kHz). To verify robustness of the aperiodic slope extraction, we additionally applied an iterative robust log-log linear regression in which frequency bins whose positive residuals exceeded 1.5 standard deviations were progressively excluded across five iterations to suppress the influence of oscillatory peaks on the aperiodic slope estimate. Results were consistent using both methods.

Asymmetry of neuronal morphology was extracted from each biophysically detailed cell type in the model. For comparison, this asymmetry was also extracted from the subthalamic nucleus neuron. The metric used to define morphological asymmetry was the effective length, defined as:

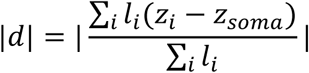

where *l*_*i*_, *Z*_*i*_ is the coordinate of its midpoint along the neuron’s principal morphological axis, and z_soma_ is the somatic coordinate along the same axis. The principal axis was defined as the first principal component of the three-dimensional point cloud formed by all dendritic segment midpoints, i.e., the axis accounting for the maximum spatial variance of the dendritic arborization. Projecting onto this axis prior to computing the effective length ensures that the metric captures the net asymmetry of dendritic mass distribution relative to the soma along the dominant morphological direction, independently of the arbitrary orientation of the reconstruction in space. Normalization by total dendritic length makes ∣d∣ independent of cell size, so that it reflects purely the degree of spatial asymmetry rather than overall morphological complexity. The absolute value is taken to ensure invariance to the sign convention of the principal axis orientation.

For each excitatory and inhibitory population within a given cortical layer, the effective length was aggregated across all reconstructed cell types assigned to that population using a weighted mean, with weights proportional to the relative numerical abundance of each cell type in the network model. This procedure yields a single population-representative morphological descriptor per layer and cell class that accounts for the heterogeneous composition of each population.

Relationships between population-averaged morphological effective length and the corresponding aperiodic PSD slopes were assessed using Pearson correlation coefficients and ordinary least-squares linear regression. Confidence intervals on regression lines were computed as 95% pointwise intervals. Statistical significance was set at p < 0.05.

Finally, to assess whether the morphology–slope relationship identified in mouse V1 generalizes to human cortical neurons, we applied the same effective length analysis to a reconstructed human layer 5 pyramidal neuron obtained from a publicly available morphological database. The effective length was computed following the identical procedure described above. The corresponding aperiodic slope was then predicted by extrapolating from the linear regression fit obtained for mouse V1 excitatory neurons (Fig. 4F).

The effects of different input rates were evaluated through a grid search across simulation parameters (Fig. 4G). The relationship between the aperiodic component and STN firing activity was assessed using the same set of simulations and the statistical methods described above, together with the excitation to inhibition imbalance, defined as:

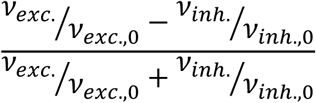

With *v*_*exc*._ And *v*_*inh.*_ the firing rate of excitatory and inhibitory inputs to STN population, *v*_*exc*.,0_ And *v*_*inh.,0*_ the values adopted in the rest of the work.

## Acknowledgements

This work was supported by the Italian Ministry of University and Research under the NRRP project “Fit4MedRob—Fit for Medical Robotics” (Grant No. PNC0000007), by Fondo Beneficenza Intesa Sanpaolo through the “GENTE” project (CUP B53C25006580007), and by the Michael J. Fox Foundation for Parkinson’s Research (MJFF) through the TuneIn project, “Retune gait freezing in Parkinson’s disease”. T. V. N. received funding from the European Union Horizon 2020 Research and Innovation Programme under Grant Agreement No., 101147319 [EBRAINS 2.0].

