## Supplementary material for "Can we trust subthalamic local field potentials? Geometric and dynamical constraints on the interpretation of extracellular recordings": Suppl. Fig.

### Supplementary Figures

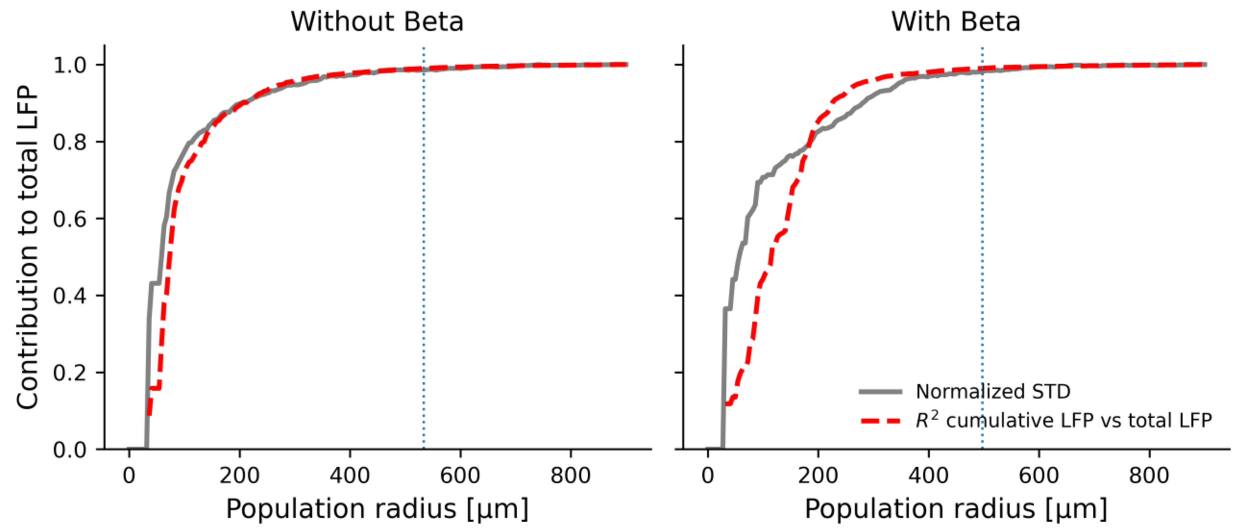


**Suppl. Fig. 1. Quality of the approximation of the total LFP from the cumulative LFP as a function of increasing radius.**

Left: Results obtained in the absence of beta oscillations. Right: Results obtained in the presence of beta oscillations. The red curve shows the standard deviation of the cumulative LFP generated within a sphere of radius *r*, normalized by the standard deviation of the cumulative LFP computed with a fixed radius of 900 µm. The gray curve shows the coefficient of determination ($R^2$) between the cumulative LFP computed within a sphere of radius *r* and the cumulative LFP computed with a fixed radius of 900 µm. The blue dashed lines indicate the radius at which the cumulative standard deviation value is 0.99 times that at 900 µm.

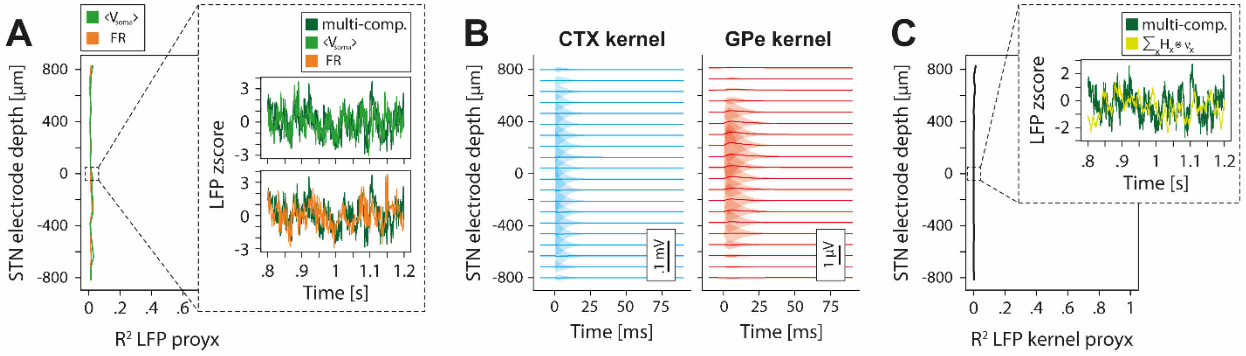


**Suppl. Fig. 2. Approximations of the LFP through population quantities and synaptic kernel:** A) Prediction accuracy of simulated STN LFPs based on the average somatic membrane potential <$V_{soma}$> and population firing rate (FR). B) Synaptic kernels at different electrode depths for cortical (CTX) and pallidal (GPe) inputs. C) Prediction accuracy of simulated STN LFPs obtained from the linear combination of the synaptic kernels.
